# Ecophysiological genomics identifies a pleiotropic locus mediating drought tolerance in sorghum

**DOI:** 10.1101/2021.12.03.471162

**Authors:** Fanna Maina, Abdou Harou, Falalou Hamidou, Geoffrey P. Morris

**Affiliations:** Department of Agronomy, Kansas State University, Manhattan, KS, USA 66506; Institut National de la Recherche Agronomique du Niger, Niamey, Niger; International Crops Research Institute for the Semi-Arid Tropics – Sahelian Center, Niamey, Niger; Department of Biology, Faculty of Sciences and Technology, Abdou Moumouni University, Niamey, Niger; Department of Soil & Crop Science, Colorado State University, Fort Collins, CO, USA 80523

**Keywords:** Genome-wide association study, Adaptation, Crop, Abiotic stress, Transpiration

## Abstract

Drought is a key constraint on plant productivity and threat to food security. Sorghum (*Sorghum bicolor* L. Moench), a global staple food and forage crop, is among the most drought-adapted cereal crops, but its adaptation is not yet well understood. This study aims to better understand the genetic basis of preflowering drought in sorghum and identify loci underlying variation in water use and yield components under drought. A panel of 219 diverse sorghum from West Africa was phenotyped for yield components and water use in an outdoor large-tube lysimeter system under well-watered (WW) versus a preflowering drought water-stressed (WS) treatment. The experimental system was validated based on characteristic drought response in international drought tolerance check genotypes and genome-wide association studies (GWAS) that mapped the major height locus at *QHT7*.*1* and *Dw3*. GWAS further identified marker trait associations (MTAs) for drought-related traits (plant height, flowering time, forage biomass, grain weight, water use) that each explained 7–70% of phenotypic variance. Most MTAs for drought-related traits correspond to loci not previously reported, but some MTA for forage biomass and grain weight under WS co-localized with staygreen post-flowering drought tolerance loci (*Stg3a* and *Stg4*). A globally common allele at S7_50055849 is associated with several yield components under drought, suggesting that it tags a major pleiotropic variant controlling assimilate partitioning to grain versus vegetative biomass. The GWAS findings revealed oligogenic variants for drought tolerance in sorghum landraces which could be used as trait predictive markers for improved drought adaptation.

## INTRODUCTION

Drought is the most common abiotic stressor limiting plant productivity globally (Barbé and Lebel, 1997). Plants species may use a variety of drought adaptive mechanisms such as escape mechanisms (e.g. early flowering to the life cycle), avoidance mechanisms (e.g. reduced plant size or transpiration), or tolerance mechanisms (e.g. reactive oxygen detoxification or osmotic adjustment) (Blum, 2004; Juenger, 2013). Still, for most plant species, the ecophysiological and genetic mechanisms underlying variation for drought adaptation in the field is not known — deciphering the mechanisms of drought adaptation remains a major challenge in plant biology and breeding (Mickelbart *et al*., 2015; Tardieu *et al*., 2018; Tuberosa, 2012). The challenge is due in part to the fundamental complexity of drought adaptation, due to pleiotropy, epistasis and genotype-environment interactions (Juenger, 2013; Tardieu *et al*., 2018), as well as the practical difficulty of studying drought stress in field conditions given year-to-year variation in rainfall (Atlin *et al*., 2017; Cooper *et al*., 2014). Genome-wide association studies in controlled environment or managed drought stress have recently been used to address this challenge, and investigate the genetic variation in drought tolerance across diverse plant germplasm (Faye, Akata, *et al*., 2021; Guo *et al*., 2018; Varshney *et al*., 2014; Wang *et al*., 2016).

Semiarid regions have experienced prolonged periods of drought that have led to severe crop yield losses in smallholder farming systems while shifts in precipitation patterns were observed in recent years (Burke *et al*., 2009). In future climates, the frequency of extreme events such as drought and floods are expected to increase in many regions (Biasutti, 2019). This is especially true in the Sahelian and Soudanian zones of Africa, large bands of semiarid grassland and savannah, respectively, where severe drought is common (Barbé and Lebel, 1997). In smallholder farming systems in the Sahel, recurrent droughts can occur throughout the growing season and limit crop production, with 10–50% yield reductions observed (Sultan *et al*., 2013; Traoré *et al*., 2011). Drought stress in the Sahel could be moderate or severe (short/intermittent and long periods of water deficit, respectively) throughout the growing season. Given such climatic variability, developing plant varieties with drought tolerance, particularly in terms of yield stability, becomes among the major priorities in breeding programs (Rapsomanikis, 2015; Walker and Alwang, 2015).

Sorghum (*Sorghum bicolor* L. Moench) is a C4 cereal crop adapted to different environmental conditions, including drought-prone areas where many other staple crops fail (National Research Council, 1996; Smith and Frederiksen, 2000). In West Africa, where dual-purpose sorghum (grain and forage) landraces are a key food security crop, the species exhibits abundant diversity across agroclimatic zones and botanical types (caudatum, durra, guinea, bicolor, and intermediates) (Bhosale *et al*., 2011; Faye, Maina, *et al*., 2021). One physiological mechanism that could contribute to drought adaptation in African landraces is staygreen post-flowering drought tolerance. During post-flowering drought, staygreen genotypes maintain their photosynthetic activity more than senescent lines due to water-saving growth and development dynamics at earlier stages (Borrell, Mullet, *et al*., 2014; Borrell, Oosterom, *et al*., 2014). Four major stay-green QTLs were reported (*Stg1–4*), which were obtained from an Ethiopian sorghum but may have a broader distribution in Africa (Borrell, Oosterom, *et al*., 2014; Faye, Akata, *et al*., 2021; Harris *et al*., 2007). For preflowering drought tolerance, a few loci have been mapped, but the underlying physiological and genetic mechanisms have not been characterized (Tuinstra *et al*., 1996; Upadhyaya *et al*., 2017a). Indeed, despite sorghum’s reputation for exceptional drought tolerance among staple crops, the physiology and genetics underlying drought adaptation in sorghum landraces is not well understood (Upadhyaya *et al*., 2017a).

To better understand the ecophysiological and genetic basis of drought adaptation in sorghum landraces, we conducted a GWAS of the drought response of diverse germplasm under managed stress in an outdoor lysimetry system. Given the differential adaptation of sorghum to semi-arid vs. sub-humid zones in West Africa (Craufurd *et al*., 1999), we hypothesized that this germplasm harbors major variants (oligogenic variation) for drought resilience traits. Further, based on studies of drought tolerance mechanisms in sorghum and other plants (Barnabás *et al*., 2008; Borrell, Oosterom, *et al*., 2014), we hypothesized that variation in plant development and water-use dynamics underlies variation in drought tolerance. Thus, our goals in this study were (i) to test these overarching hypotheses on the genetic and physiological basis of drought adaptation in sorghum landraces and (ii) to generate specific hypotheses on the genetic loci and physiological processes involved. Our findings suggest that major variants for drought tolerance exist in African sorghum landraces, and that some affect multiple drought-related traits pleiotropically.

## MATERIAL AND METHODS

### Plant materials

The West African sorghum association panel (WASAP), consisting of landraces and breeding lines, was previously assembled from sorghum breeding programs in four countries (Mali, Niger, Senegal, and Togo), genotyped, and phenotyped for agro-morphological traits (Faye, Maina, *et al*., 2021). In this study, we used a subset of 219 genotypes from the WASAP, hereafter referred to as WASAP_Lysi (Table S1; Data S1). WASAP_Lysi is a curated subset of the WASAP, selected based on the genotypic data to capture most of the genetic diversity existing in West Africa (Fig. 1A-B, Fig. S1A-B) and based on phenotype data (Faye, Maina, *et al*., 2021) to remove highly photoperiod sensitive genotypes that would confound drought phenotyping under common-garden conditions (Tuberosa, 2012). In addition, two breeding lines that have been widely used for drought tolerance studies in the U.S. and Australia were included as checks for preflowering drought tolerance (Tx7000) and postflowering drought tolerance (BTx642) (Borrell, Mullet, *et al*., 2014; Borrell, Oosterom, *et al*., 2014; Harris *et al*., 2007; Tuinstra *et al*., 1996).

**Fig. 1.**
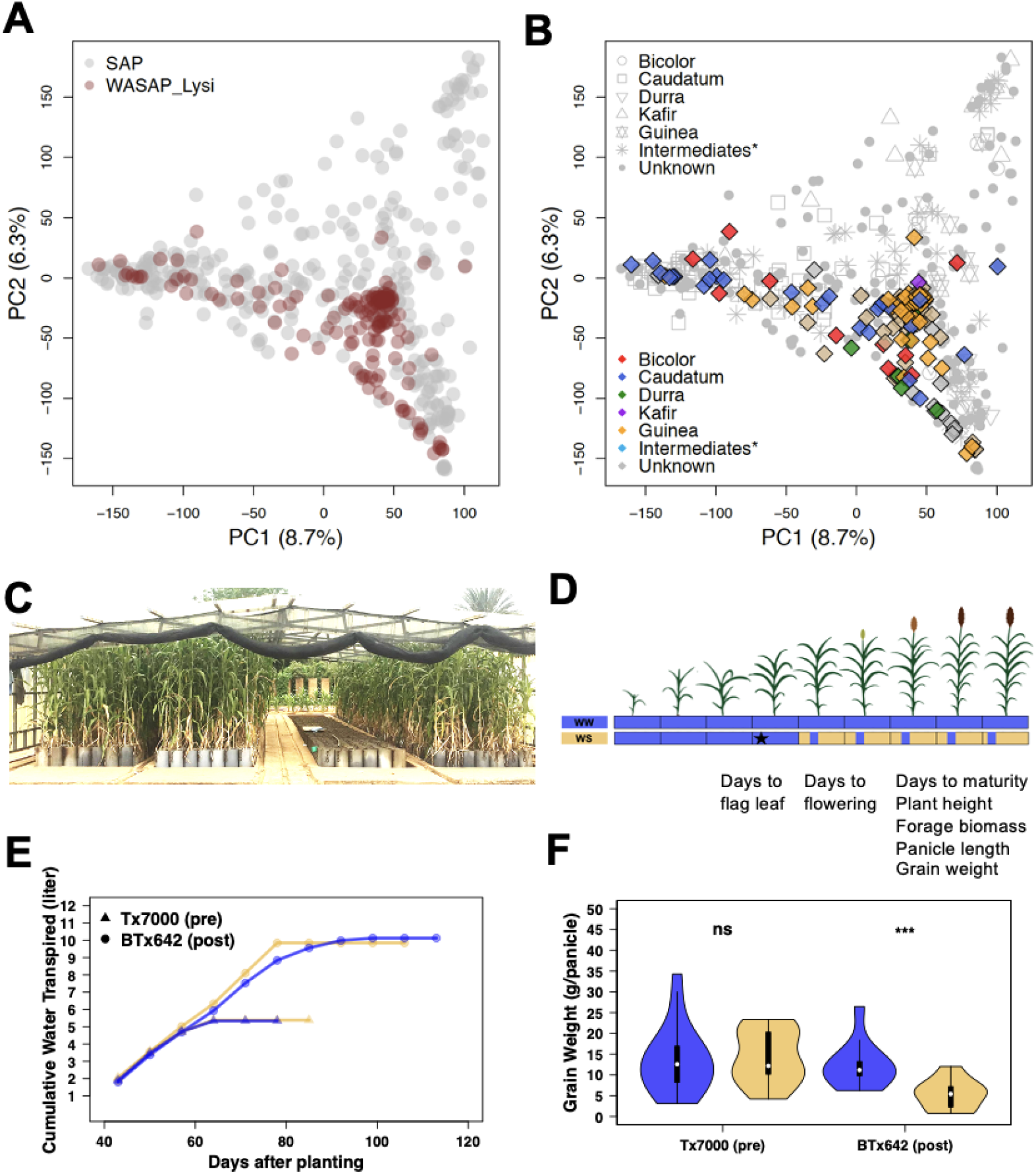
Experimental system for managed drought phenotyping of diverse sorghum. (A) Principal coordinate analysis (PCA) of WASAP_Lysi (maroon) compared to a global sorghum association panel (SAP) (in grey) and (B) with botanical type noted. (C) Lysimeter facility at ICRISAT Sadoré-Niger, with plants grown in large tubes (1.30 m × 0.25 m) during the normal cropping season under a retractable rain-out shelter. (D) Schematic representation of well-watered (WW, blue) and intermittent preflowering water-stressed (WS, yellow) treatments, with phenotype collected listed below. To account for phenology differences among the genotypes, WS was imposed from flag leaf appearance for each individual at different times (represented by the star). (E) Cumulative water use for Tx7000 and BTx642 and (F) grain weight (g/panicle) in WW and WS treatments for international drought tolerance check lines, Tx7000 (preflowering drought tolerant; “pre”) and BTx642 (postflowering drought tolerant; “post”).

### Genomic diversity analyses

Genotypic data used was generated using the genotyping-by-sequencing (GBS) method with the *Ape*KI restriction enzyme (Faye, Maina, *et al*., 2021; Hu *et al*., 2019). To characterize the genetic diversity of WASAP_Lysi relative to the global sorghum diversity (sorghum association panel; SAP) (Fig. 1A-B) and the West African sorghum diversity (as represented in USDA-GRIN genebank; WAGRIN) (Fig. S1), we analyzed sequence reads obtained from previous studies (Faye *et al*., 2019; Lasky *et al*., 2015; Maina *et al*., 2018; Morris *et al*., 2013; Olatoye *et al*., 2018). The reference genome BTx623 version 3.1 (McCormick *et al*., 2018; Paterson, 2013) was used to align sequencing data using Burrows-Wheeler Alignment (Li and Durbin, 2010), and the SNPs were discovered using TASSEL 5 GBS pipeline (Glaubitz *et al*., 2014). Missing SNPs were imputed in Beagle v1.4 (Browning and Browning, 2009). SNPs were filtered at MAF > 0.01 and only biallelic sites were kept. A total of 219 WASAP_Lysi genotypes, 1527 West African genotypes, and 342 SAP genotypes were further analyzed (Fig. 1A-B). To characterize the WASAP_Lysi genetic diversity relative to the SAP and the WAGRIN, principal component analysis (PCA) was performed in R using the *SNPRelate* package (Zheng *et al*., 2012). First, the variance components of each genotype in the WASAP_Lysi using the training set from SAP with ∼90,000 biallelic SNPs present in both collections (WASAP_Lysi and SAP) was predicted. Next, the variance components of WASAP_Lysi genotypes were predicted using WAGRIN as a training set with ∼138,000 biallelic SNPs present in both collections (WASAP_Lysi and WAGRIN).

### Lysimeter phenotyping system

Experiments were conducted in a lysimeter system at the ICRISAT Sahelian Center, Niger (Sadoré, 13.15°N, 2.18°E) during the normal cropping season from June to November, in 2017 and 2018. The rainy season at this location is June to October. The lysimeter system, equipped with a rainout shelter, consists of evaluating the physiological characteristics of crops under managed experiments in large PVC tubes (Falalou *et al*., 2018; Vadez *et al*., 2008, 2013). Tubes of 1.30 m in height and 0.25 m in diameter were placed in trenches (Halilou *et al*., 2015) and filled with field soil from a farm in Sadoré station (Fig. 1C). Each tube was fertilized with 6 g of 15–15–15 (N–P_2_O_5_–K_2_O) following a recommended standard fertilization (Ministère de l’agriculture du Niger, 2012). The soil surface was covered with transparent plastic bags and 350 g of polyethylene beads in each lysimeter tube to limit soil evaporation.

### Water treatments

The experiment was conducted in a split-unit design with two treatments, well-watered (WW) and water-stressed (WS), and with three replicates. The experimental unit was each lysimeter tube. Three seeds were planted in each tube, and two plants were left in the tube a week after emergence. Two weeks after planting, one individual plant in each tube was kept for phenotyping. The two genotypes with known drought stress response, Tx7000 and BTx642 were highly replicated in both treatments (*n* = 14) during the second year of the experiment (Borrell, Mullet, *et al*., 2014). In the well-watered treatment (WW), genotypes were irrigated at 80% field capacity until physiological maturity. In the WS treatment, genotypes were irrigated at 80% field capacity, as in the WW, until fully flag leaf appearance for each genotype, then intermittent drought stress was applied (Fig. 1D). The WS treatment consisted of skipping irrigation upon the observation of a wilting point on the last three leaves before relieving the water stress with one liter of water.

### Trait phenotyping

Plant height was measured from the base of the stem to the tip of the last fully emerged leaf each week. Days to flowering was defined as the number of days until anther appearance on each plant. Yield components include final forage biomass, on a per plant basis (vegetative biomass in g/plant), and grain weight, on per panicle basis (g/panicle). Beginning six weeks after planting, lysimetric tubes (one individual plant in each tube) were weighed weekly using a hanging scale (Mettler Toledo; 20 g accuracy) to estimate weekly water use on a per plant basis. Total water transpired after weighing was estimated weekly as the total water added until physiological maturity of each lysimeter tube.

### Statistical analysis of phenotypes

Analyses were performed in the R program (R Core Team, 2013). To estimate the variance components across years within a treatment, the *lme4* package was used (Bates *et al*., 2015):

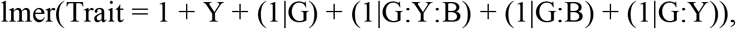

where G is genotype, Y is year, and B is replicate block. Year is treated as a fixed effect. Best linear unbiased predictions (BLUPs) were then estimated for genome-wide association studies. Broad-sense heritability across years within the same treatment was estimated as:

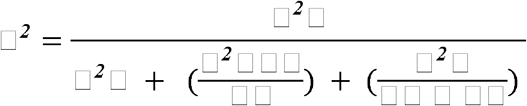

where □^*2*^□ is the genotypic variance, □^*2*^ □ is the residual variance, *nY* is the number of years, *nR* is the number of replicate blocks, and □^*2*^□□□is the genotype-by-year interaction.

ANOVA was performed to assess whether the sources of variation were significant for the evaluated traits between treatments for each country and botanical type on a mean basis. Drought response was calculated as the difference between well-watered and water-stressed treatments on a mean basis for the two years. The broad-sense heritability was estimated using the estimate of genetic variance (□^*2*^□) and the residual variance (□^*2*^ □).

### Genome-wide association studies

To identify marker-trait associations, we used 73,730 genome-wide SNP markers in WASAP_Lysi after filtering for minor allele frequency (MAF = 0.05). Next, we conducted GWAS for yield components in both treatments separately for WW, WS, and the difference between WW and WS. BLUPs were estimated with the linear model above mentioned and carried out GWAS using the general linear model (GLM) and mixed linear model (MLM) and multilocus mixed linear model (MLMM) (Lipka *et al*., 2012; Segura *et al*., 2012) for plant height, days to flowering, forage biomass, and grain weight for WW and WS treatment across two years. The difference between WW and WS phenotypic values was estimated for each genotype as phenotypes. We used the Bonferroni correction threshold (conservative threshold, α = 0.05, *p*-value < 10^−7^) to identify significant markers associated with the phenotypes. To further identify outlier SNPs (marker-trait associations, MTAs) associated with each trait, the top 0.01% lowest *p*-values in the MLMM model were selected.

To determine whether MTAs co-localized with previously identified and putative genes, a list of *a priori* candidate genes was generated from a literature review (File S2). This list (*n* = 67) of candidate genes includes their genomic positions in sorghum and orthologs of rice and maize for plant height, maturity, grain yield, biomass, and drought tolerance. The sorghum QTL Atlas was also used to determine the co-localization of MTAs with previously identified QTLs from the literature (Mace *et al*., 2018). Putative orthologs were identified based on the highest similarity in the “Protein Homologs” in Phytozome 13 and expression data was from GeneAtlas v1 Tissue Sample in Phytozome 13 (Goodstein *et al*., 2012). Geographic distribution of alleles was based on existing GBS resources for georeferenced sorghum landraces (Hu *et al*., 2019; Lasky *et al*., 2015).

## RESULTS

### Genetic structure of the diversity panel

To first characterize the genetic structure of our experimental panel (WASAP_Lysi) with respect to global sorghum diversity, we analyzed 90,148 genome-wide SNPs. Taking the SAP as representative of the global diversity, the first principal and second principal component explained 8.7% and 6.3% of the total variation, respectively (Fig. 1A-B). When the WASAP_Lysi was projected on the principal components of SAP (Fig. 1A), the WASAP_Lysi genotypes clustered with the caudatum, durra, and guinea types in SAP on the first two PCs (Fig. 1B). To characterize the genetic structure of WASAP_Lysi with respect to West African sorghum diversity, we conducted PCA on 138,000 genome-wide SNPs. Consistent with our observations in the SAP, PC1, PC2, and PC3 distinguish major groups of caudatum, durra-caudatum, and guinea respectively (Fig. S1A-B). Since the majority of WASAP_Lysi have not been classified into botanical types morphologically, we used genotyping data to classify accessions into botanical types. We inferred botanical types from admixture (*K* = 3, cross-validation error = 0.62) using membership probability > 0.6 to assign WASAP_Lysi genotype to a botanical type for further analyses (Fig. 1B).

### Phenotypic variation and drought response

To validate the quality of the managed drought lysimeter phenotyping we first characterized drought response phenotypes for two international drought tolerance checks, Tx7000, the preflowering drought tolerance check, and BTx642, the preflowering drought susceptible check. Tx7000 had less water transpired compared to BTx642 in both treatments (Fig. 1E). There were no significant differences in total water transpired for either BTx642 and Tx7000 between treatments (*p* = 0.6 and *p* = 0.9, respectively; Fig. 1E). Comparing WS versus WW phenotypes for grain yield and harvest index, there is no reduction for Tx7000, but a substantial reduction for BTx642 (*p* < 0.001) (Fig. 1F, Fig. S2). The maximum water use in both genotypes was reached two weeks after flowering. In both genotypes, the drought response on flowering time, plant height, and forage biomass was negligible (Fig. S2). Days to flowering was slightly delayed in the WS treatment for Tx7000 (+3 days; *p* < 0.08) but not for BTx642 (*p* < 0.2) (Fig. S2). The performance of the checks was similar in both years (Fig. S3).

Next, we characterized phenotypic variation for ecophysiological traits across the WASAP_Lysi diversity panel. Overall, a significant genotype effect was observed for all morphological and phenological traits, in both WW and WS treatments (*p* < 0.05; Table 1). Under both WW and WS, broad-sense heritability estimates were high (0.75–0.97), with the highest heritability for days to flowering in WW (Table 1). The heritabilities of the drought response (WW-WS) of forage biomass and grain weight were high (0.86 and 0.50, respectively), while the heritabilities of drought response of plant height and days to flowering were low (0.05 and 0.09, respectively) (Table 1). Within each botanical type and country of origin, a significant reduction of grain weight was observed in WS compared to WW, as would be expected (Fig. S4).

**Table 1.** Means, variance components, and heritability for BLUPs of five traits under well-watered and water-stressed treatment and their drought response.

| Trait<sup>a</sup> | Well-watered (WW) |  |  |  |  | Water-stressed (WS) |  |  |  |  | Response (WW-WS) |  |  |  |
| --- | --- | --- | --- | --- | --- | --- | --- | --- | --- | --- | --- | --- | --- | --- |
|  | Mean | SD | G | GxY | H<sup>2</sup> | Mean | SD | G | GxY | H<sup>2</sup> | Mean | SD | G | H<sup>2</sup> |
| PH | 255 | 46 | 48**<br>* | 25 | 0.96 | 252 | 45 | 49*** | 8*** | 0.96 | 2 | 22.5 | 0 | 0.05 |
| DF | 91 | 23 | 24**<br>* | 1 | 0.96 | 92 | 4 | 25*** | 3 | 0.97 | -1 | 9 | 40 | 0.09 |
| FB | 105 | 22 | 27**<br>* | 16*** | 0.77 | 92 | 23 | 26*** | 14*** | 0.83 | 12 | 14.5 | 12*** | 0.86 |
| GW | 12.6 | 6 | 55**<br>* | 17* | 0.75 | 8 | 5 | 40*** | 7* | 0.79 | 4.5 | 3 | 6.7*** | 0.50 |

<sup>a</sup> Traits: PH: Plant height (cm); FB: Forage biomass (g); DF: Days to flowering (days); ; GW: Grain weight per panicle (g).
SD: standard deviation; G: Genotype; GxY: Genotype-by-Year interaction; H<sup>2</sup>: Broad-sense heritability. Significance at $p < 0.05$ , 0.01, and 0.001 for two-way ANOVA is note with \*, \*\*, and \*\*\*, respectively. G and GxY terms without asterix did not pass the significance threshold.

Water use in the WASAP_Lysi showed significant genotypic variation (*p* < 10^−15^). In all of the botanical types, cumulative water use was reduced significantly (*p* < 0.01) under WS vs. WW treatment (Fig. 2A-B). On average, caudatum types had less cumulative water use compared to durra-caudatum and guinea types in both treatments (Fig. 2A-B). The majority (80%) of the genotypes showed a reduction of cumulative water use in WS compared to WW, as much as 15 liters in some genotypes (Fig. 2C). Heritability of water use was moderate (0.4–0.6) in the first five weeks and high thereafter (up to ∼0.8 by physiological maturity) under both WW and WS treatments (Fig. 2D).

**Fig. 2.**
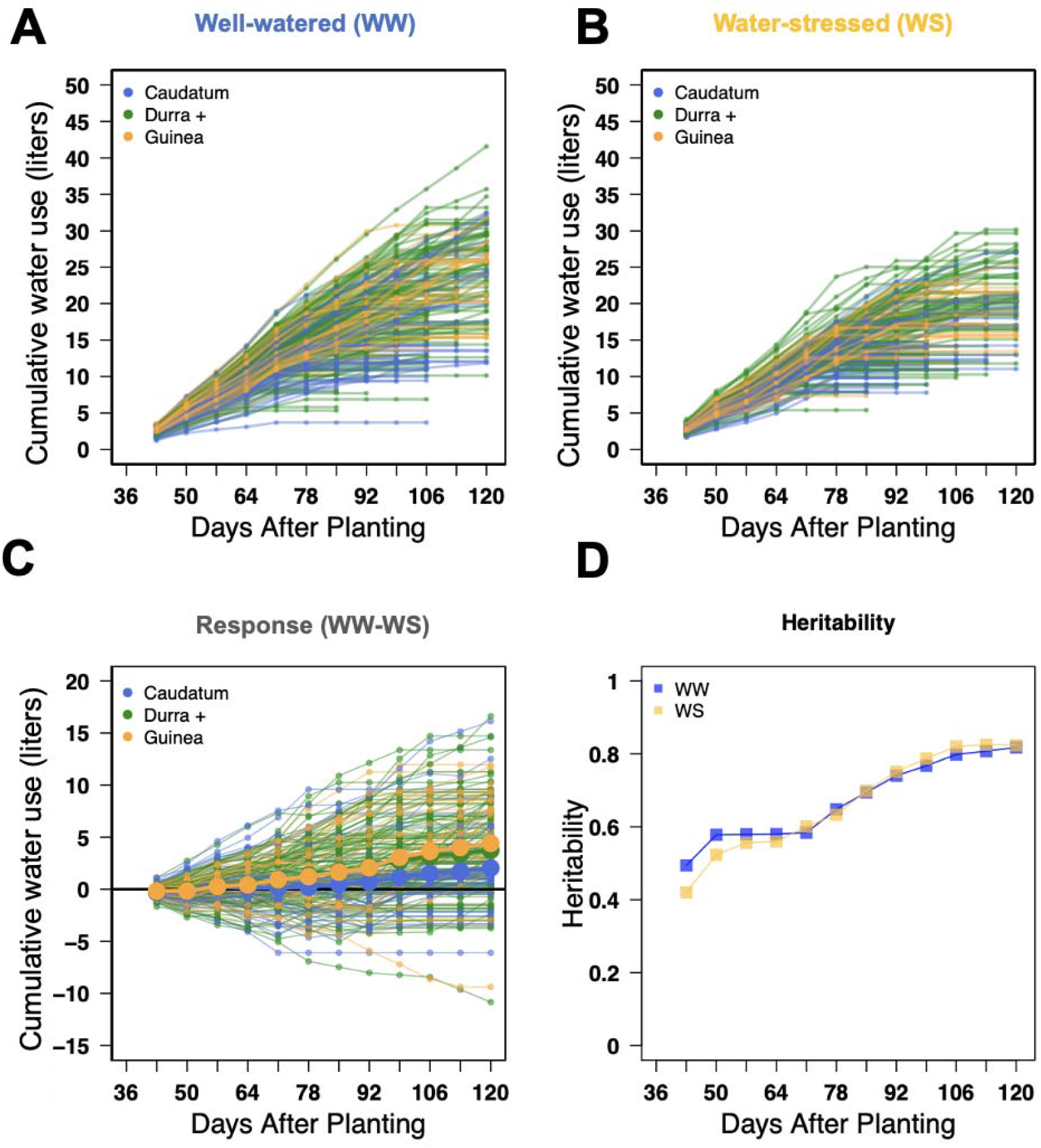
Genetic variation for water use under drought, by botanical type and genotype. Cumulative water use in well-watered (WW; A) and water-stressed (WS; B) treatments, and drought response for water use (WS-WS) (C). Genotypes were color-coded by botanical type. Averages of each botanical type are represented in thicker lines color-coded by type, with large solid circles representing the mean for each botanical type. (D) Broad-sense heritability estimates for cumulative water use by time point under WW (blue) and WS (yellow) treatments.

### Marker-trait associations for plant height and flowering time and their drought response

To validate our experimental system for GWAS, we first identified MTAs for plant height and flowering time for each year individually using a standard approach of GLM and MLM GWAS (Fig. S5-S6). MTAs for plant height on chromosomes 4, 6, 7, and 8 explained between 11% and 25% of the total phenotypic variance for the 2017 data. Among the associations, S7_59412395 (MAF = 0.05) explained 19% of the variation and co-localized with the classical dwarfing gene *Dw3* (∼0.4 Mb) (Fig. S5). In 2018 data, 19 MTAs were identified on chromosomes 6, 7, and 10 (Fig. S5). The phenotypic variation explained by the MTAs ranged from 12% to 23%. With MLM, ten MTAs on chromosomes 7 and 10 in both years explained 12–19% of variation. For flowering time, three MTAs were identified via GLM on chromosome 6 (S6_50696803; MAF = 0.13, S6_50716126; MAF = 0.16, and S6_50716244; MAF = 0.16) for 2017 data, whereas no significant MTA were identified in 2018 data (Fig. S6). These flowering time MTAs did not co-localize with the *a priori* candidate genes for maturity on chromosome 6 (*Ma6, Ma1, SbCN4, abph1, zfl1*; Data S2).

To further validate the experiment and gain insights on these drought-related traits, we conducted MLMM GWAS on plant height and flowering time using across-year BLUPs, for each treatment (Fig. 3A-F). (Note, MLMM are designed to identify individual MTA at a QTL, rather than broad association peaks of multiple MTA typical of GLM and MLM). Under WW treatment, a few significant associations were identified, two for plant height and one for days to flowering, respectively (Fig. 3A, 3D). For plant height, the top association was S7_59412395 (*p* < 10^−20^; MAF = 0.05), roughly co-localized with the classical dwarfing gene *Dw3* (∼0.4 Mb) and closely colocalized with the top MLMM MTA from a field-based GWAS of the WASAP in Senegal (S7_59400476; 11 kb away) (Faye, Maina, *et al*., 2021). The second most significant MTA was S6_2900751 (*p* < 10^−7^; MAF = 0.09), which is within the QTL interval *QHGHT6*.*4* (2.37–3.6 Mb) (Takai *et al*., 2012). The proportion of the phenotypic variance explained (PVE) by the MTAs is 17% (Table 2).

**Table 2.** Marker-trait associations from MLMM above the Bonferroni threshold (α = 0.05)

| **Trait** | **Treatment<sup>a</sup>** | **SNP** | **$-\log(p)$** | **MAF<sup>b</sup>** | **Allele (Mean phenotype)** | **PVE<sup>c</sup> (%)** |
| --- | --- | --- | --- | --- | --- | --- |
| Flowering | WW | S4_64398335 | 8.6 | 0.07 | G (88), C (145), S (77) | 46 |
| Flowering | WW-WS | S5_52090779 | 7.5 | 0.07 | T (-1), C (46), Y (0) | 70 |
| Flowering | WS | S3_19589652 | 7.4 | 0.13 | C (93), Y (106), T (94) | 7 |
| Flowering | WS | S4_14172212 | 6.9 | 0.08 | A (89), G (138), R (126) | 39 |
| Grain Weight | WS | S7_50055849 | 9.8 | 0.14 | T (7), A (14), W (5) | 27 |
| Grain Weight | WS | S6_46923493 | 6.7 | 0.05 | C (7.4), A (13), M (5.8) | 12 |
| Grain Weight | WS | S6_18075344 | 6.4 | 0.18 | G (6.7), A (11), R (8) | 33 |
| Plant Height | WS | S7_59459123 | 23 | 0.06 | T (263), A (140), W (292) | 12 |
| Plant Height | WW | S7_59412395 | 19 | 0.05 | G (267), A (125), R (237) | 48 |
| Plant Height | WW-WS | S4_149131 | 7.7 | 0.04 | G (3), C (-35), S (19) | 23 |
| Plant Height | WS | S7_50055849 | 7.4 | 0.14 | T (269), A (209), W (261) | 39 |
| Plant Height | WS | S9_56534065 | 6.6 | 0.07 | T (259), G (251), K (231) | 6 |
| Plant Height | WW | S6_2900751 | 6.5 | 0.09 | G (265), A (206), R (234) | 5 |

<sup>a</sup> WW: well-watered; WS: water-stressed; WW-WS: drought response.
<sup>b</sup> Minor allele frequency.
<sup>c</sup> Percentage of phenotypic variance explained based on linear model.

**Fig. 3.**
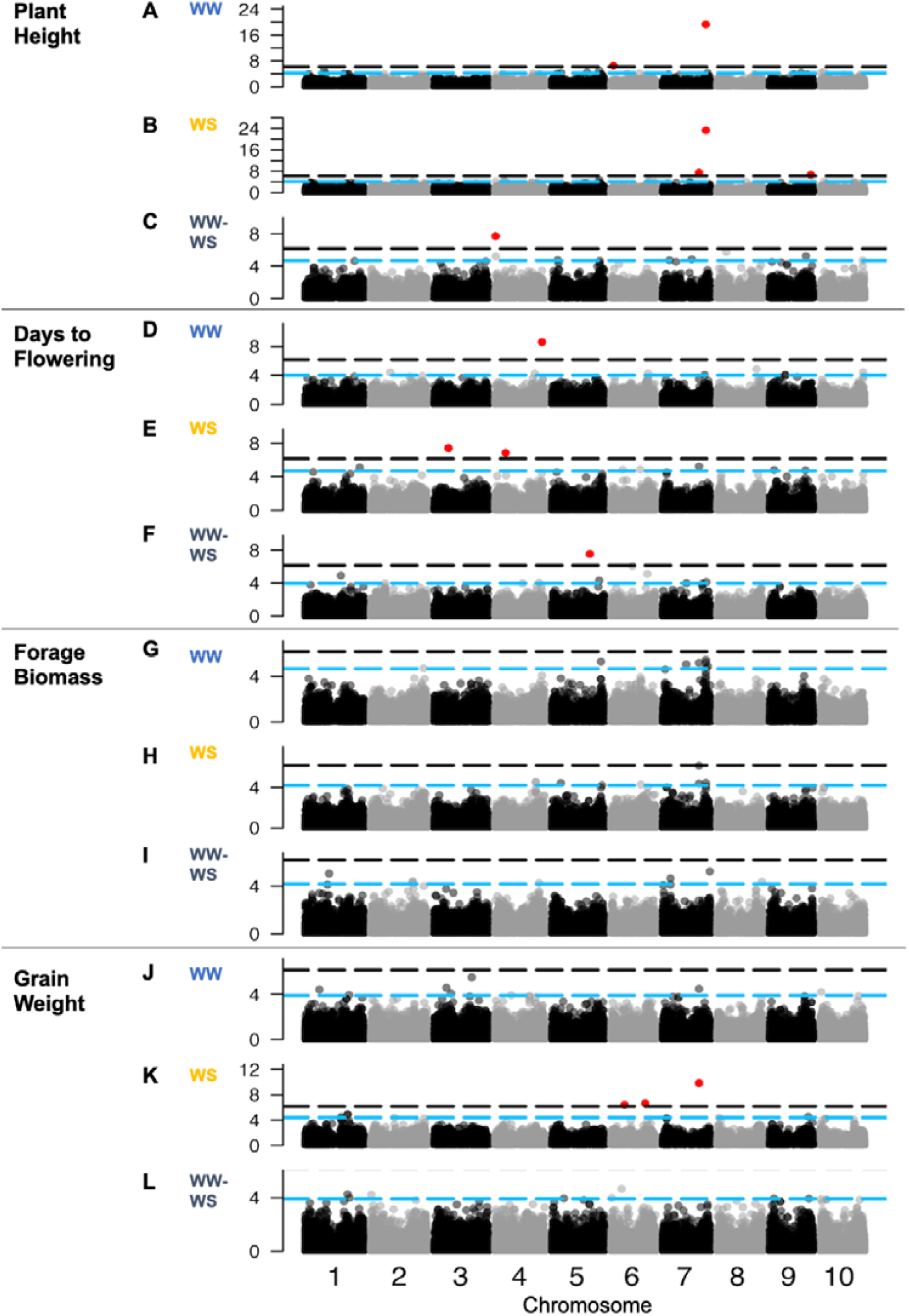
Genome-wide association study for plant traits and their drought response. Multi-locus mixed model associations for traits in well-watered treatment (WW; A, D, G, J), water-stressed treatment (WS; B, E, H, K), and the drought response of the traits in WW versus WS treatments (WW-WS; C, F, I, L). Black dashed lines represent the conservative threshold of α = 0.05 with a Bonferroni correction (with red points indicating marker-trait associations above the threshold), while the blue line represents 0.01% lowest *p*-values.

For flowering time, S4_64398335 (MAF = 0.07; *p <* 10^−9^; 46% PVE) was found within the QTL region *QDTFL4*.*25* (61.02–65.08 Mb) (Mocoeur *et al*., 2015). Under WS treatment, there were eight MTAs across all traits (Fig. 3B, 3E). Three MTAs for plant height were observed, with the top association at S7_59459123 near *Dw3* (MAF = 0.06), along with associations at S7_50055849 (MAF = 0.14) and S9_56534065 (MAF = 0.07). The latter MTA overlaps with QHGHT9.45 (56.1–57.8Mb) for plant height (Fig. 3B) (Felderhoff *et al*., 2012). There were two MTAs for days to flowering under WS conditions (Fig. 3E), on chromosome 3 (S3_19589652; MAF = 0.13) and 4 (S4_14172212; MAF = 0.11). The majority of Malian and Togolese genotypes are later flowering, associated with the minor allele (C) at S4_64398335. Togolese genotypes (*n* = 18) with the G allele at S4_14172212 flower later under WS treatment.

To identify loci that may underlie variation in drought response for flowering time and height, we conducted MLMM on the difference between WW and WS (WW-WS). One MTA for drought response of plant height was identified above the conservative threshold at S4_149131 (MAF = 0.04) (Fig. 3C). Another flowering time MTA was found above the conservative threshold on chromosome 5 (S5_52090779; MAF = 0.07; *p <* 10^−7^; 70% of the phenotypic variance explained) (Fig. 3F, Table 2), which co-localized with QDTFL5.8 (9 Mb region) for days to flowering (Mace *et al*., 2013). Among the top 0.01%, MTAs for drought response of flowering time (Data S3), S5_64932515 (MAF = 0.05, Fig. 3F) co-localized with flowering time locus QDTFL5.10 (0.8 Mb) (Srinivas *et al*., 2009).

### Marker-trait associations for forage and grain yield components and their drought response

Next, to identify loci that may underlie variation for yield under preflowering drought, we conducted MLMM for forage and grain yield components under WW and WS, and their drought response (Fig. 3G-L). Under WW treatment, no associations for forage biomass and grain weight per panicle were above the conservative threshold (Fig. 3G, 3J). To identify MTAs that fall below the conservative threshold but may still be of interest, we investigated MTAs with the slowest 0.01% *p*-values for each trait (Data S3), and noted cases where they are colocalized with previously identified QTL. Under WW treatment, the top MTA for forage biomass was S7_59459123 (MAF = 0.06; *p* < 10^−5^), which was also a plant height MTA and co-localized with the *Dw3* gene (290 kb away). Another MTA for forage biomass, S2_72712959 (MAF = 0.19; *p* < 10^−5^), co-localized with QTDBM2.4 (total dry biomass, 70–76 Mbp) (Mocoeur *et al*., 2015). For forage biomass under WS treatment (Fig. 3H), the top MTA was at S7_50055849 and the fourth-highest was the same MTA (S7_59459123) observed under WW treatment (Data S3).

For the drought response of forage biomass, the top MTA is on chromosome 7 (S7_64933026; *p* < 10^−5^; Fig. 3I). Note, this MTA is not colocalized with the plant height MTAs near *Dw3* and *QHT7*.*1*. In addition, three MTAs were observed (S2_57620549, S2_57664163, and S2_57663973) spanning a 43 kb region within staygreen locus *Stg3a* (57.1–61.8 Mbp) (Fig. 3I, Data S3, Fig. S7). The minor alleles associated with lower forage biomass under drought were at high frequency in the panel (MAF = 0.42, 0.38, and 0.39, respectively).

For grain weight per panicle under WS, three associations were found, on two chromosome 6 (S6_18075344, MAF = 0.18; S6_46923493, MAF = 0.05) and one on chromosome 7 (S7_50055849; MAF = 0.14) (Fig. 3K). For the drought response of grain weight, an MTA was identified S6_18075344, which was also identified under WS treatment (Fig. 3L; Table 2). The average values for the genotypes with the G allele (mean = 4.9 g) at S6_18075344 (which included BTx642) were greater than the ones with the A allele (mean = 4.6 g) (which included Tx7000) (Table 2). Genotypes originated from Mali (*n* = 7), Niger (*n* = 21), and Senegal (*n* = 3) have the same preflowering drought tolerance-associated alleles as Tx7000. By contrast, at the second most associated marker (S6_46923493) (Fig. 3L), the minor allele (A) associated with moderate reduction in grain weight under WS, is carried only by Nigerien genotypes (*n* = 12). One MTA for drought response of grain weight (S5_16199662; MAF = 0.09, *p* > 10^−4^) co-localized with *Stg4* (Fig. 3L). Genotypes with the non-reference allele at this marker (A, *n* = 31) had less of a reduction in grain weight under drought than genotypes with the reference allele (G, *n* = 153).

### Marker-trait associations for water use and its drought response

To identify loci that may underlie variation in water use dynamic and play a role in drought avoidance, the water use of each genotype under both WW and WS treatment was measured weekly over twelve weeks and MTAs were identified. Using MLMM for water use at each time point, a total of 62 MTAs were identified. Several MTAs appear at two or three consecutive time points (e.g. S2_63494925, S8_51257440, S8_570367, and S9_32820789) among the top 0.01% of the lowest *p*-values (Fig. 4A). At 85 days after planting, a MTA was observed at S2_63675212 (MAF = 0.23), which colocalized with *Stg3b* (62.7–64.6 Mb).

**Fig. 4.**
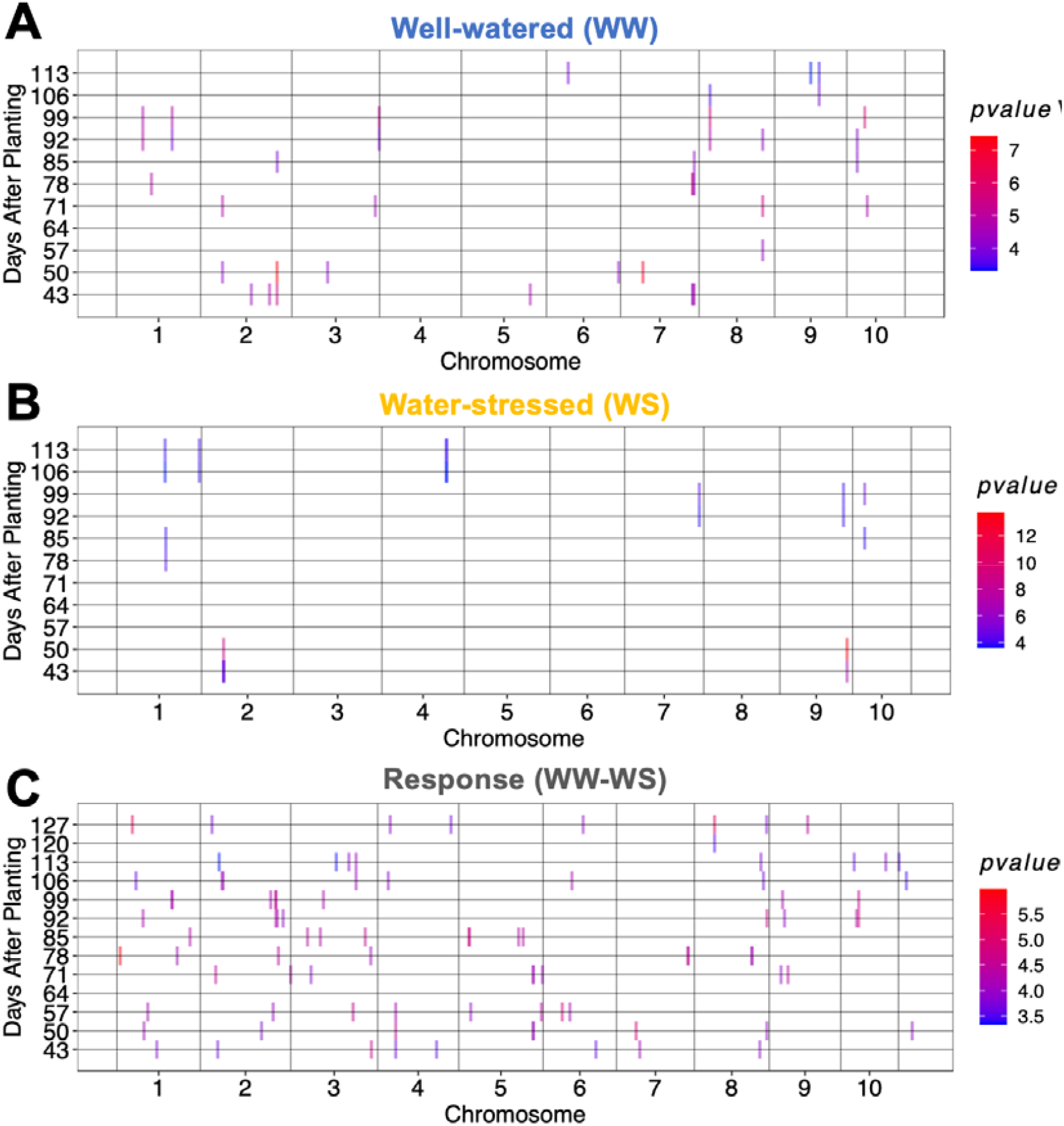
Genome-wide association study of water use and its drought response. Multi-locus mixed linear model genome-wide association studies for weekly water use under (A) well-watered treatment, (B) water-stressed treatment, and (C) for difference between well-watered and water-stressed treatment. The *x*-axis represents the genomic position of each marker, and the *y*-axis represents the days after planting. Vertical bars (color-coded by -log of the *p*-values) represent the genomic position of marker-trait associations at each of the 12 time points.

For water use under WS treatment, 12 MTA were identified. While some of the MTAs were observed at a single time point, the majority of MTAs (84%) were shared across two or more consecutive time points (Fig. 4B). The strongest association for water use under WS was on chromosome 9 (S9_54114067, *p* < 10^−14^, MAF= 0.36) at 50 days after planting. For the drought response, a total of 89 MTAs among the top 0.01% of the lowest *p-*values were observed across time points (Fig. 4C). Some drought response MTAs appear at two or three consecutive time points. For instance, the MTA at S4_7506983 is observed at consecutive time points 43, 50, and 57 days after planting. Another MTA at S2_63345899 (*p* < 10^−5^) was found within the staygreen region *Stg3b* (62.7–64.6 Mb) at two consecutive time points after 90 days.

### Putative pleiotropic QTL and their geographic distribution

To identify loci that may be pleiotropically influencing multiple drought-related traits we cataloged associations that were shared across multiple traits (Fig. 5A). Shared associations (n = 5) on chromosomes 4, 5, and 7 were observed for multiple traits. For instance, variation at S7_59459123 (MAF = 0.06) is associated with variation in plant height in both water treatments (Fig. 5A). Most notably, allelic variation at S7_50055849 (MAF = 0.14) is associated with variation of several traits we characterized: grain weight and forage biomass under WS, plant height under WS, days to flowering under WS, forage biomass (WW, WS), and grain weight (WW, WS) (Fig. 5B). The minor allele (A) is associated with shorter stature, earlier flowering, less forage biomass, and greater grain weight per panicle (*p* = 0.005).

**Fig. 5.**
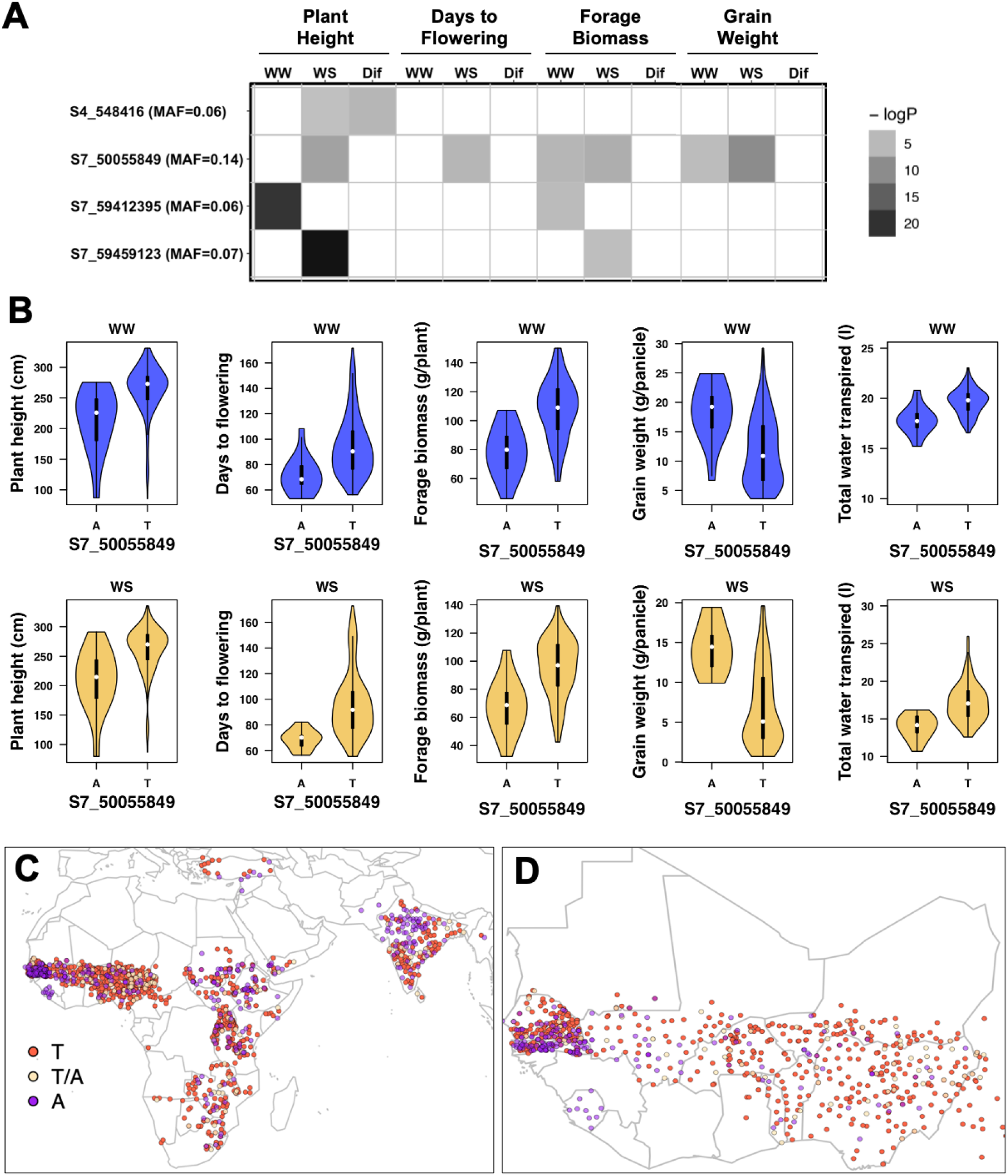
Shared marker-trait associations across traits identify putative pleiotropic loci. (A) Heatmap for -log(*p*-values) of shared marker-trait associations for plant height, days to flowering, forage biomass, grain weight, and total water use in well-watered (WW) and water-stressed (WS) treatments. (B) Phenotypes for accessions carrying the A (minor allele) or T (major allele) at SNP S7_50055849, a putative pleiotropic locus. Distribution of alleles at S7_50055849 in georeferenced accessions (C) across Africa and Asia, and (D) in West Africa.

Within the WASAP_Lysi, accessions carrying the minor allele at S7_50055849 (A; MAF = 0.14) originated from Niger (*n* = 21), Mali (*n* = 4), and Senegal (*n* = 2), while none of the Togolese germplasm (mostly from the guinean agroclimatic zone) carries the minor allele. To characterize the global origin and prevalence of the putative variant, we mapped the global distribution of alleles at S7_50055849 across georeferenced landraces (Fig. 5C-D). The minor allele is common (21%) across Africa and in India (Fig. 5C). Within West Africa, the minor allele is distributed at high frequency in southern Senegal and Sierra Leone compared to Niger, Nigeria, Mali, and Burkina Faso (Fig. 5D).

## DISCUSSION

### Using ecophysiological genomics to understand drought adaptation in sorghum

Ecophysiological genomics, including quantitative genomic analysis of ecophysiological traits, could contribute to a better understanding of the genetic basis of drought tolerance. Sorghum landraces harbor extensive diversity of drought response phenotypes (Haussmann *et al*., 1998; Lasky *et al*., 2015; Upadhyaya *et al*., 2017b), but it is not known what aspects of the phenotypic response contribute to drought resilience. The goals of this study were to test some hypotheses on the genetic architecture and physiological basis of drought adaptation in African sorghum landraces and to generate specific hypotheses on loci that could underlie variation in drought adaptive traits. In sub-Saharan Africa, drought can occur throughout the growing season, from germination to grain filling stages (Biasutti, 2019). We used managed stress experiments to repeatably investigate stress scenarios that are important in the target population of the environment (TPE) but are difficult to study due to unpredictable occurrence (Cooper *et al*., 2014). We focused on preflowering drought because (1) it is less well-studied and understood than postflowering drought (Borrell, Oosterom, *et al*., 2014) and (2) it is tractable to precisely apply water deficit at the same phenological stage, even for diverse germplasm with flowering time variation. We studied a large regional landrace diversity panel that captures much of the global diversity of sorghum (Fig. 1A).

High-quality phenotypic data is essential for dissecting complex traits, such as those underlying drought adaptation (Cobb *et al*., 2013; Mackay *et al*., 2009). For stress response traits particularly, quality phenotyping requires extensive controls using genotypes with known stress responses to validate the experimental system (Tuberosa, 2012). Several lines of evidence suggest that high-quality managed stress phenotyping was obtained in this study, such that resulting findings are relevant to phenotypic response of sorghum to natural field-based drought. First, treatment-by-year effects were minor for both WW and WS (*p* = 0.99 and 0.94, respectively), suggesting that unintended sources of variation were minimized (Fig. S2-S3). Second, genotype-by-environment interaction was observed between international drought tolerance check genotypes, in the expected direction based on their known preflowering drought reaction. Finally, the lower reduction in grain weight for Tx7000 than BTx642 (Fig. 1F, Fig. S2) confirmed that the stress imposed in our study is comparable to natural preflowering drought stress (Kebede *et al*., 2001; Tuinstra *et al*., 1996). Still, further research will be required to test the hypothesis that the lysimeter managed drought stress (Fig. 1) provides a good proxy for drought in the TPE, for instance, by confirming that managed stress phenotypes (lysimeter system) are positively correlated to those in the TPE (multi-environmental trials) (Cooper *et al*., 1995).

### Evidence of oligogenic variation for drought tolerance

A key hypothesis we aimed to test in this study is that sorghum landraces harbor a few major variants for preflowering drought tolerance (oligogenic hypothesis). If this hypothesis is true, then the effect of individual drought tolerance variants can be characterized, and these variants could be deployed in marker-assisted selection (Bernardo, 2008). However, if variation for preflowering drought tolerance is due to many variants of small effect (polygenic hypothesis), then characterizing individual variants will be unnecessary and infeasible, and phenotypic or genomic selection would be required. As would be predicted under the oligogenic hypothesis, a modest number MTAs (n = 13) were identified across traits and treatments (Table 2), a few co-localized with previously identified QTLs and others not previously described (Data S3). However, given that this is an association study it is not possible to eliminate the hypothesis that the novel MTAs include spurious associations (Vilhjálmsson and Nordborg, 2013). To confirm the hypotheses on genetic variants, it would be necessary to confirm phenotypic differences in drought-related traits in controlled genetic backgrounds, such as recombinant or near-isogenic lines (Bergelson and Roux, 2010).

Flowering time is a drought-related trait because early flowering can contribute to drought escape, while plant height is a drought-related trait because reduced biomass can contribute to drought avoidance (water saving) (Blum *et al*., 1989; Tuberosa, 2012). Substantial flowering time variation exists in West African sorghum (Bhosale *et al*., 2011; Upadhyaya *et al*., 2017b) that could contribute to drought escape, and we observed strong MTAs that could be contributors (Fig 3A-C; Data S3). The flowering time MTAs did not co-localize with *a priori* candidate genes for flowering time, suggesting that variation at known genes (e.g. *Maturity* loci, orthologs of maize or rice flowering genes; Data S3) does not underlie the variation we observed, and that previously unreported genes may be involved. This contrasts previous field-based studies of the WASAP that observed flowering time associations near known genes such as *Maturity6/SbGHD7, SbCN8, SbCN12*, and *SbZFL1* (Faye, Akata, *et al*., 2021; Faye, Maina, *et al*., 2021). However, in this study we selected genotypes with little to no photoperiod response (based on the previous common-garden experiments) to reduce confounding effects of flowering time variation on drought phenotyping (Tuberosa, 2012). Thus, the lack of MTA at known genes could be explained by the panel design, which excluded major photoperiodic flowering time variants. For plant height, by contrast, several MTAs were found between the two major genes controlling plant height in sorghum *QHT7*.*1* and *Dw3* (Fig. S5), which matches the field-based WASAP study (Faye, Maina, *et al*., 2021).

Among the components of drought tolerance, yield stability is particularly important in smallholder cropping systems given the severe consequences of crop failure in this context (Rapsomanikis, 2015). MTAs for the drought response of yield components (grain weight and forage biomass) are candidate loci for preflowering drought tolerance since they reflect yield stability under drought. Several MTAs with forage biomass (S2_72712959, S2_57620549, S2_57664163, and S2_57663973) and grain weight (S7_64933026, S6_46923493, S5_16199662) co-localized with previously identified QTLs for the same traits (Data S3) (Mace *et al*., 2018). Some of the top MTAs for the drought response of forage biomass (S2_57620549, S2_57664163, and S2_57663973) and grain weight (S5_16199662) (Fig. 3I-L) co-localized with *Stg3a* and *Stg4*, respectively. This finding is consistent with the hypothesis that staygreen alleles, which delay senescence under post-flowering drought in semi-dwarf grain sorghums (Borrell, Oosterom, *et al*., 2014), also contribute to drought tolerance in West African landraces (Faye, Akata, *et al*., 2021).

### A major pleiotropic variant mediating water use and drought tolerance?

The most notable association in the study is the putatively pleiotropic MTA at S7_50055849 (Fig. 5A), which is the top association for both grain weight and forage biomass under drought (Fig. 3H-K, Fig. 5A) as well as a leading association for flowering time (third highest) and plant height (second highest) under drought (Data S3). This SNP is also associated with grain and forage yield under WW, but in these cases the association is marginal (*p* _≈_ 10^−5^) and it is not the top association. One hypothesis to explain these findings is that a major variant exists near S7_50055849 that mediates partitioning of assimilates to grain versus vegetative biomass under drought, along with other aspects of plant growth and development (Blum, 1998; Ongom *et al*., 2016; Tuberosa, 2012). Under this assimilate partitioning hypothesis, the variant would pleiotropically affect multiple yield and drought-related traits as downstream physiological effects.

Given that S7_50055849 is associated with flowering time and height, an alternate hypothesis would be that a nearby maturity variant is delaying flowering, thus increasing forage biomass and decreasing grain yield. However, under this delayed maturity hypothesis, one would expect that the top associations for grain weight would be the top flowering time associations (e.g. S4_64398335, S3_19589652, etc.), which was not the case (Fig. 3; Data S2). Therefore, the shared associations at S7_50055849 for yield components and flowering time are unlikely to be due to delayed maturity, but more likely to be due to a variant mediating assimilate partitioning across the plant that in turn affects maturity.

Consistent with the hypothesis that the MTA at S7_50055849 represents a novel loci mediating assimilate partitioning, the MTA did not co-localize with any known maturity or height genes (Data S2). One interesting *post hoc* candidate gene near S7_50055849 is Sobic.007G115400 (second-nearest gene model, 43 kb away), an uncharacterized gene that is moderately expressed (0.6–5.5 FPKM) in stem tissues and lowly expressed in other tissues. While the putative ortholog of this gene in maize (GRMZM2G136353) and rice (Os08g28870) have not been functionally characterized, the putative ortholog in Arabidopsis (AT1G28440) is *HAESA-LIKE 1* (*HSL1*). In Arabidopsis, *HSL1* is a peptide hormone receptor that negatively regulates stomatal development (Qian *et al*., 2018). Thus, one gene-level hypothesis for the MTA at S7_50055849 is that a variant at Sobic.007G115400 affects stomatal development, and in turn grain and forage yield under drought. Validation and fine-mapping of the QTL using near-isogenic lines would be necessary to test this mechanistic hypothesis.

If there is a major pleiotropic variant near S7_50055849 as suggested by the multi-trait GWAS (Fig. 3, Fig. 5A-B), the geographic analysis of alleles at this SNP (Fig. 5C-D) suggests that the variant is widespread in landraces across both East and West Africa and South Asia. If this hypothesis is true, future genome-wide mapping studies of drought-related traits in diverse East African and South Asian germplasm should identify MTA in the same genomic region. A recent GWAS for field-based managed drought stress using the full WASAP did not identify a preflowering drought tolerance MTA at S7_50055849 but did identify one at S7_50507851, 450 kb upstream (Faye, Akata, *et al*., 2021).

### Conclusion

In this study we used an ecophysiological genomics approach, with managed drought stress phenotyping and GWAS, to investigate the physiology and genetics of preflowering drought adaptation in diverse sorghum landraces. We found abundant phenotypic variation for drought-related traits, including drought response of yield components and water use, and evidence of oligogenic trait variation, with several strong MTAs, including a putative pleiotropic drought tolerance variant near S7_50055849. If confirmed in field environments and controlled genetic backgrounds, the alleles associated with higher yield under drought or favorable drought response of yield components could be the basis for trait-predictive markers used in breeding for climate-resilience and future genetic dissection of drought adaptation (Tuberosa, 2012; Varshney *et al*., 2014).

## ACKNOWLEDGMENTS

This paper is made possible by the support of the American People provided to the Feed the Future Innovation Lab for Collaborative Research on Sorghum and Millet through the United States Agency for International Development (USAID) under Cooperative Agreement No. AID-OAA-A-13-00047. The contents are the sole responsibility of the authors and do not necessarily reflect the views of USAID or the United States Government. We are grateful to the physiology lab technicians at ICRISAT Sadoré in Niger. The authors declare they have no conflict of interest.

---

**Fig. S1.**
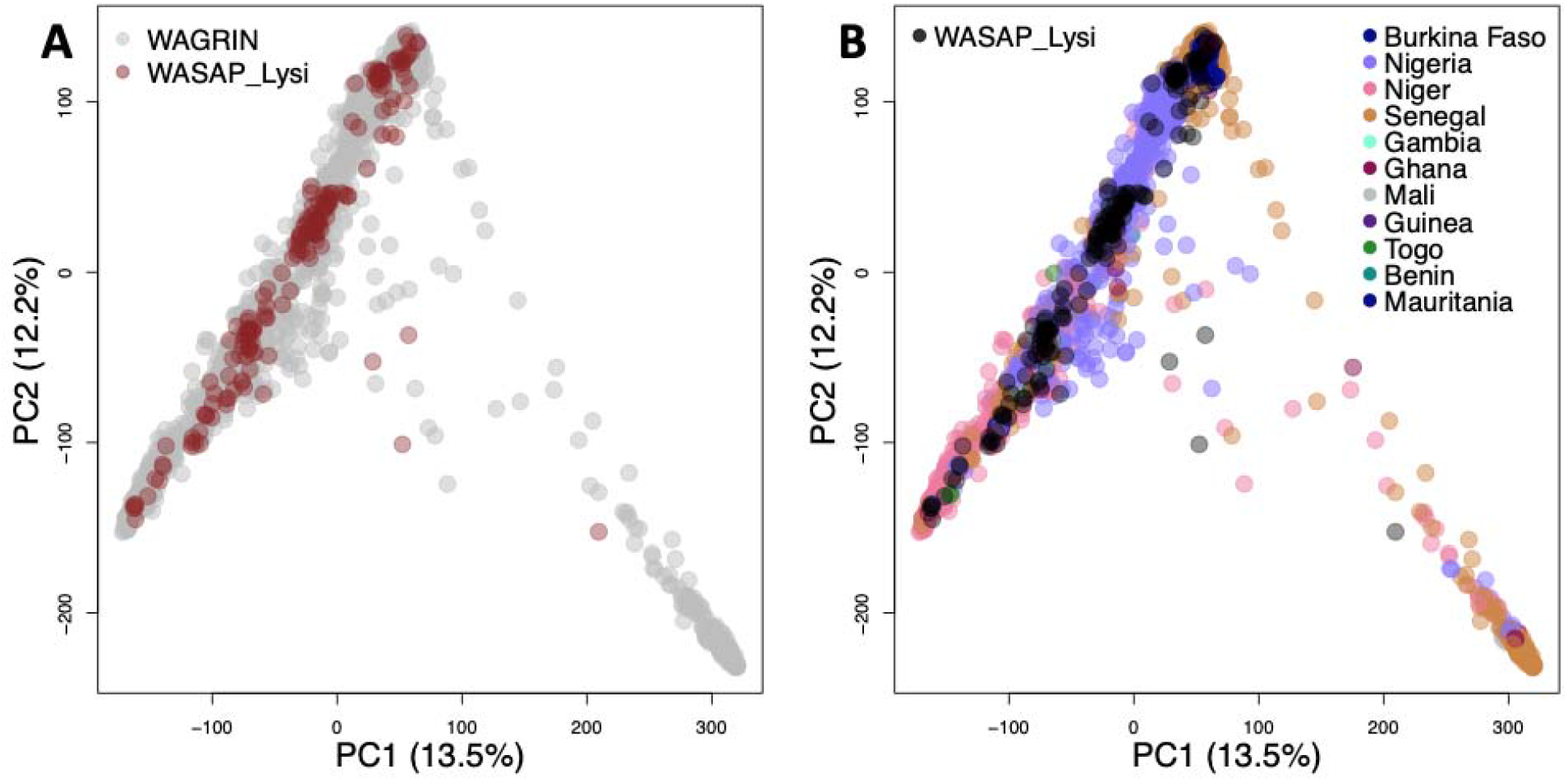
Principal component analysis (PCA) of genome-wide nucleotide polymorphism. (A) PCA of WASAP_Lysi relative to the West African genotypes of USDA-GRIN. (B) PCA of WASAP_Lysi relative to the West African genotype color-coded by country of origin.

**Fig. S2.**
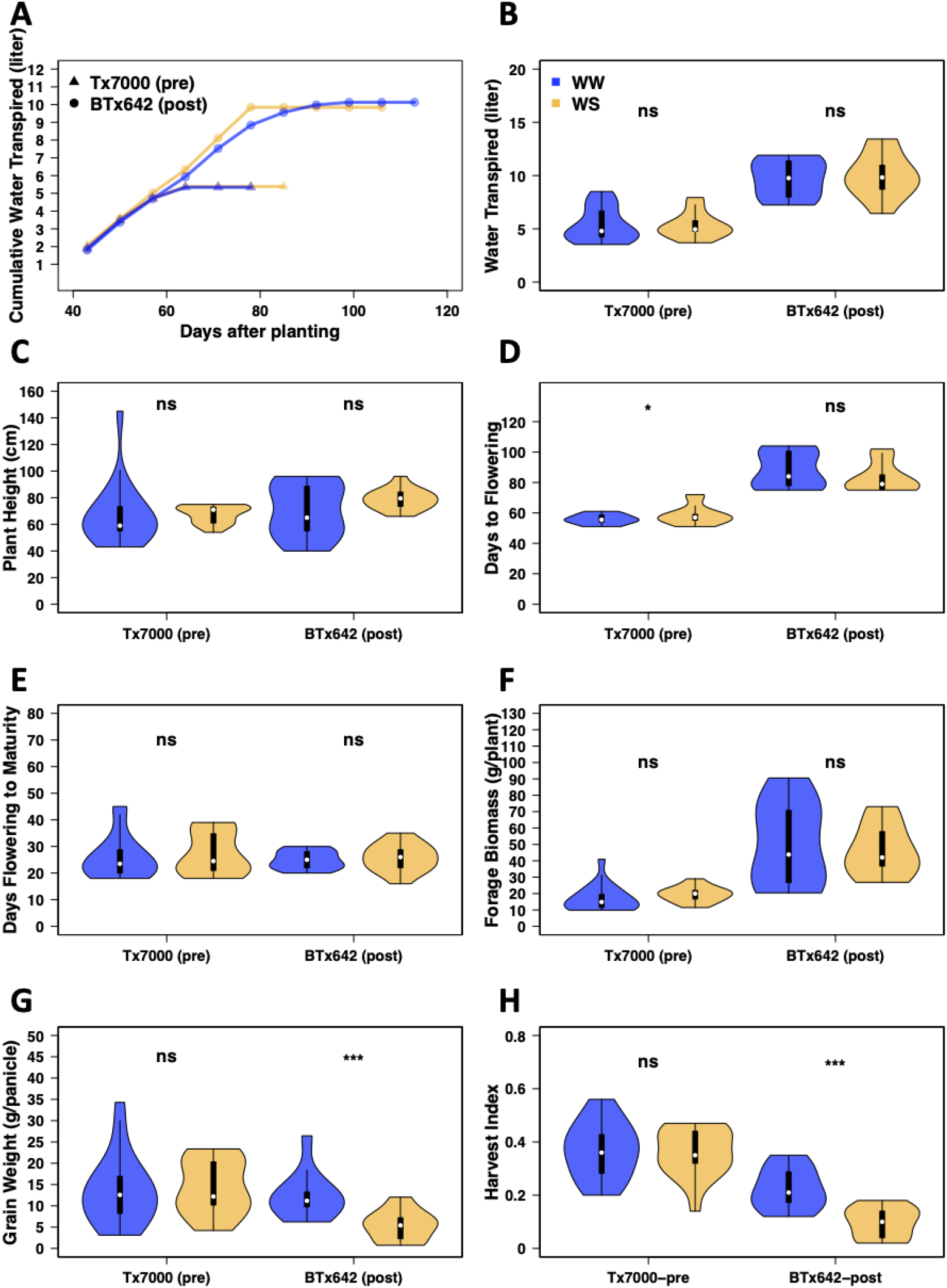
Drought response phenotypes of international drought tolerance check genotypes establish the validity of the lysimeter system. Cumulative water transpired until physiological maturity in well-watered (WW, blue violin plots) and water-stressed treatments (WS, orange violin plots) for Tx7000 (preflowering drought tolerance check; “pre”) and BTx642 (post-flowering drought tolerance check; “post”) in 2018 (*n* = 14). Asterix *, **, and ***, represent *p*-values less than α < 0.05, 0.01, and 0.001, respectively ns: not significantly different, *p* > 0.05.

**Fig. S3.**
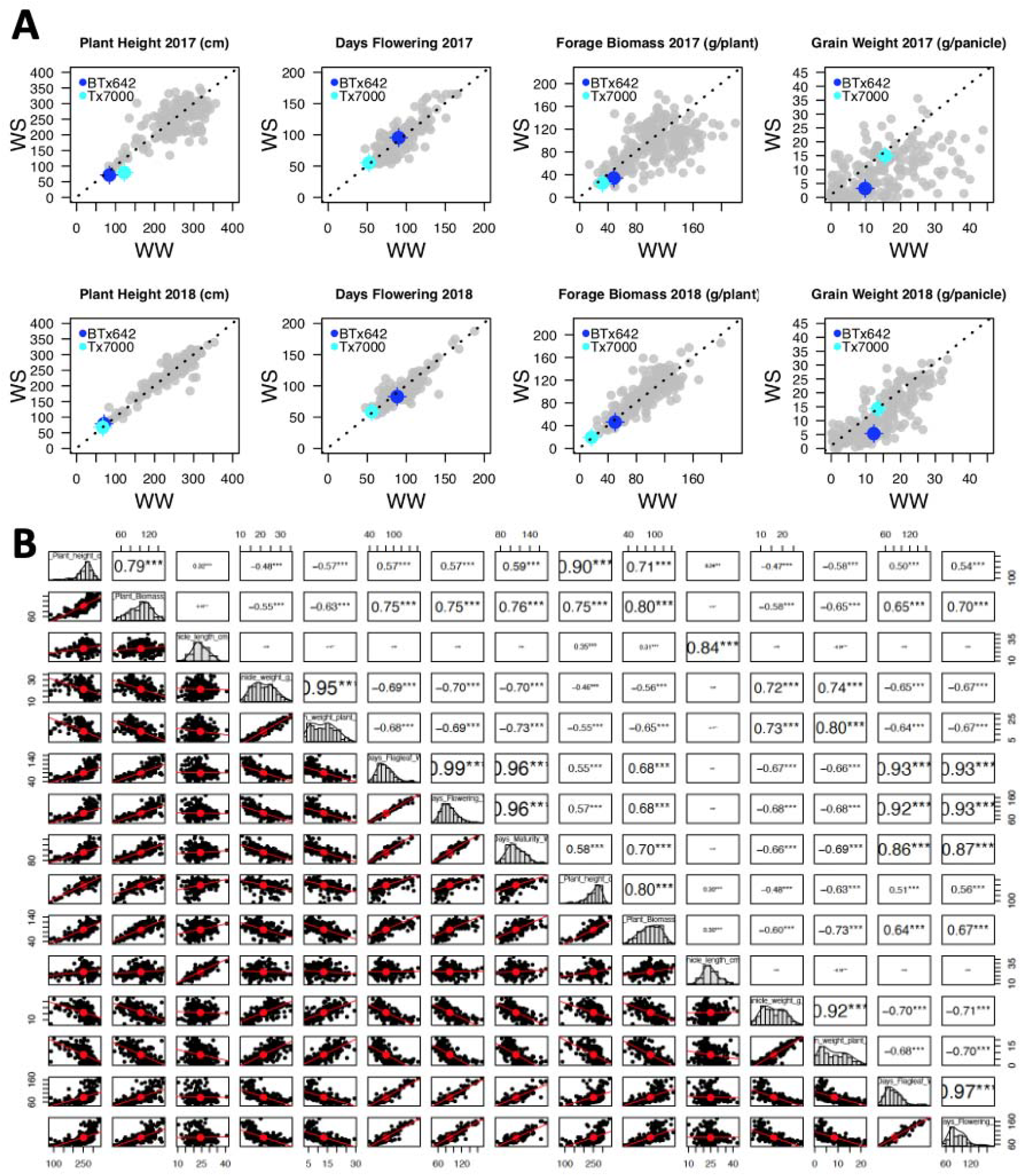
Correlations of phenotypes across years and traits. (A) Correlation across years and (B) among traits.

**Fig. S4:**
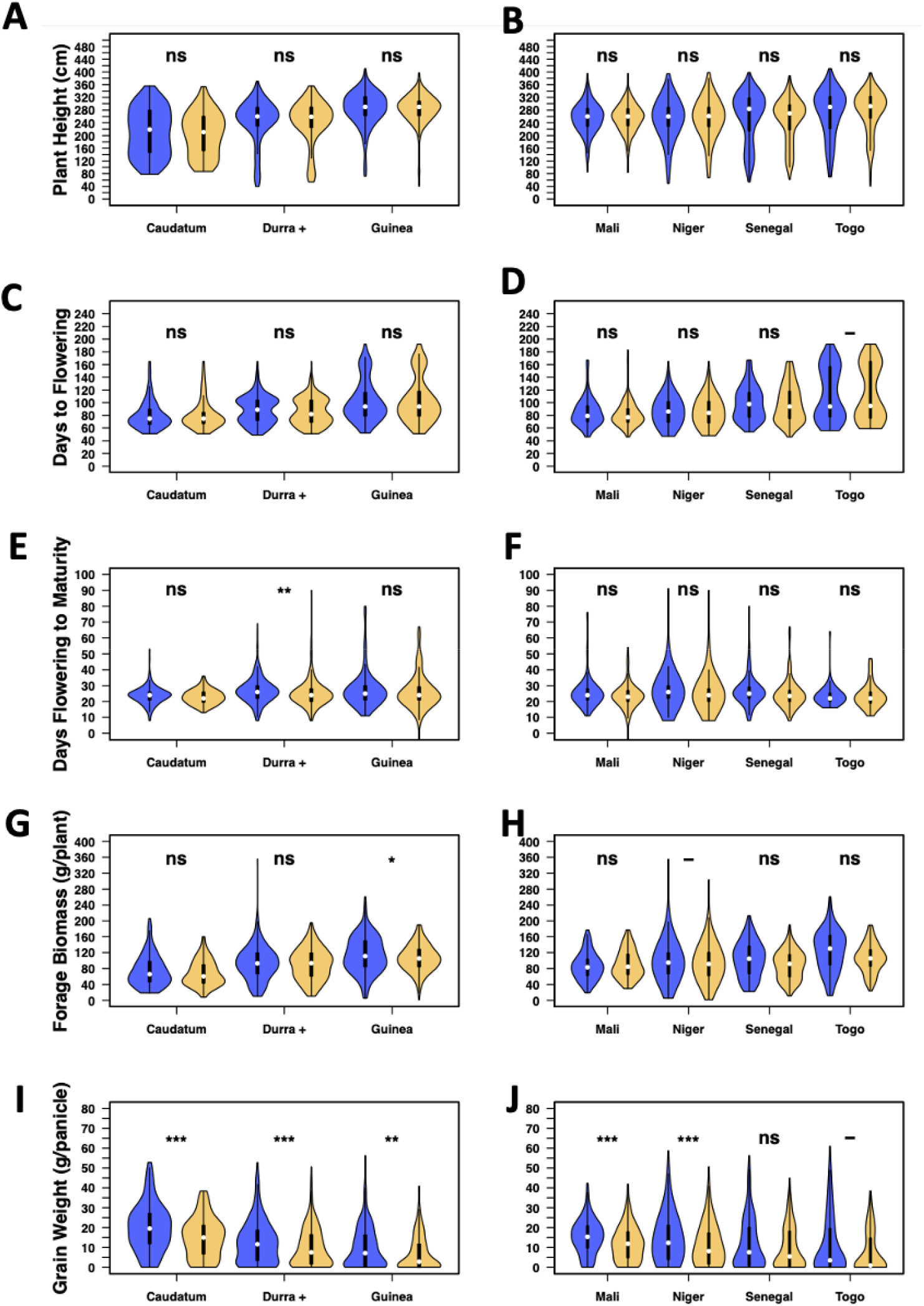
Drought response for plant height, days to flowering, Days flowering to maturity, forage biomass, and grain weight for botanical types and country of origin. Botanical types according to genotypic classification: Caudatum (n=28), Durra+ (Durra and durra-caudatum) (n=74), Guinea (n=50). Country of Origin: Mali (n=31), Niger (n=116), Senegal (n=37), Togo (n=28). Asterix *, **, and *** represent significance at *p* < 0.05, 0.01, and 0.001, respectively; ns: not significant.

**Fig. S5.**
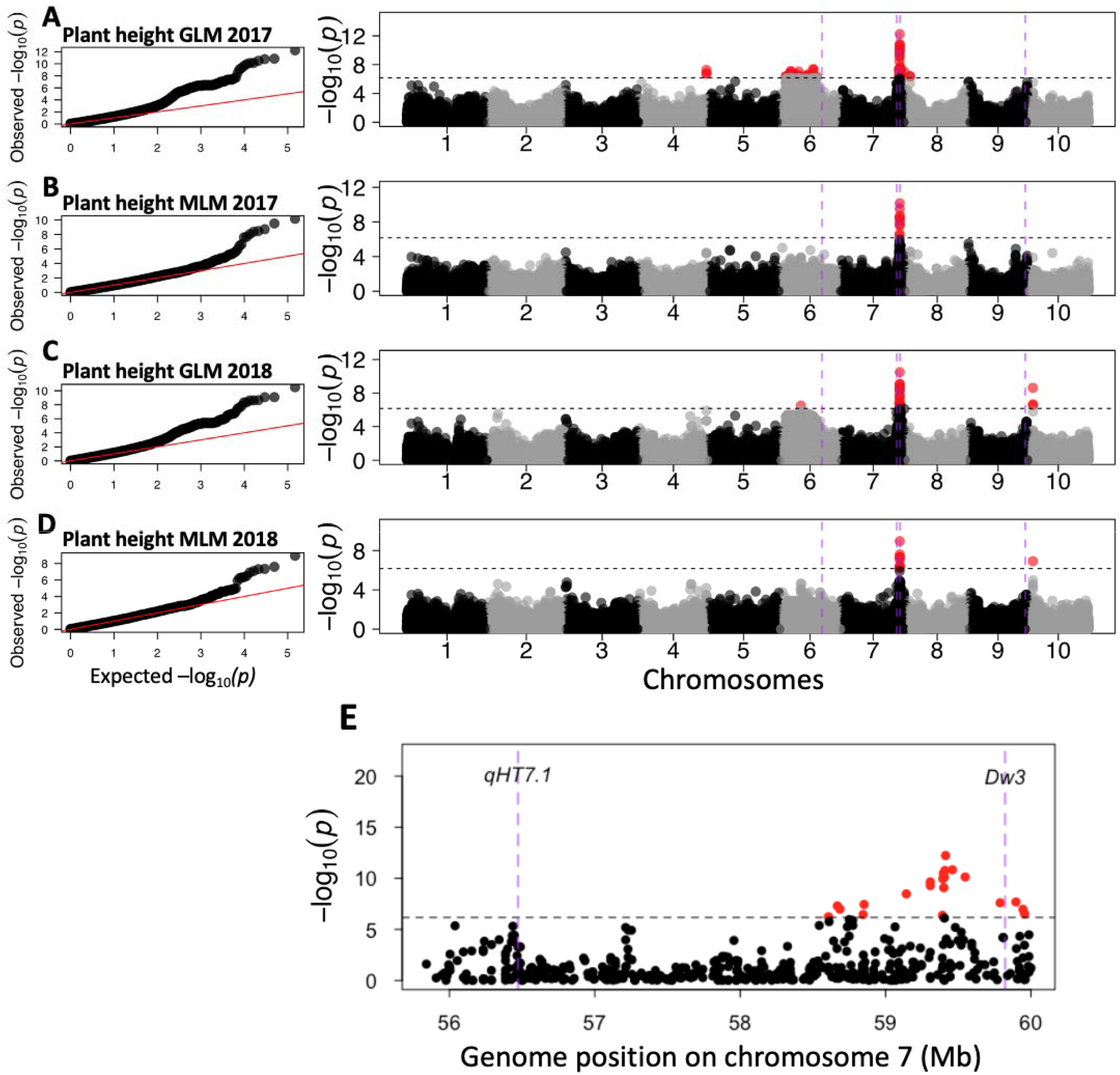
Genome-wide association studies of plant height in well-watered treatment. (A-B) Quantile-quantile and Manhattan plots in 2017 using the general linear model (GLM) (A) and mixed linear model (MLM) (B). (C-D) Quantile-quantile and Manhattan plots in 2018 using general linear model (GLM) (C) and mixed linear model (MLM) (D). (E) Zoomed-in Manhattan plot on chromosome 7 for GLM model in 2017 highlighting *qHT7*.*1* and *Dw3* genomic regions. Vertical dashed lines (purple) represent *a priori* candidate genes for plant height *Dw2* (chromosome 6), *qHT7*.*1* (chromosome 7), *Dw3* (chromosome 7), and *Dw1* (chromosome 9). Horizontal dashed lines represent the threshold at Bonferroni correction (α = 0.05).

**Fig. S6.**
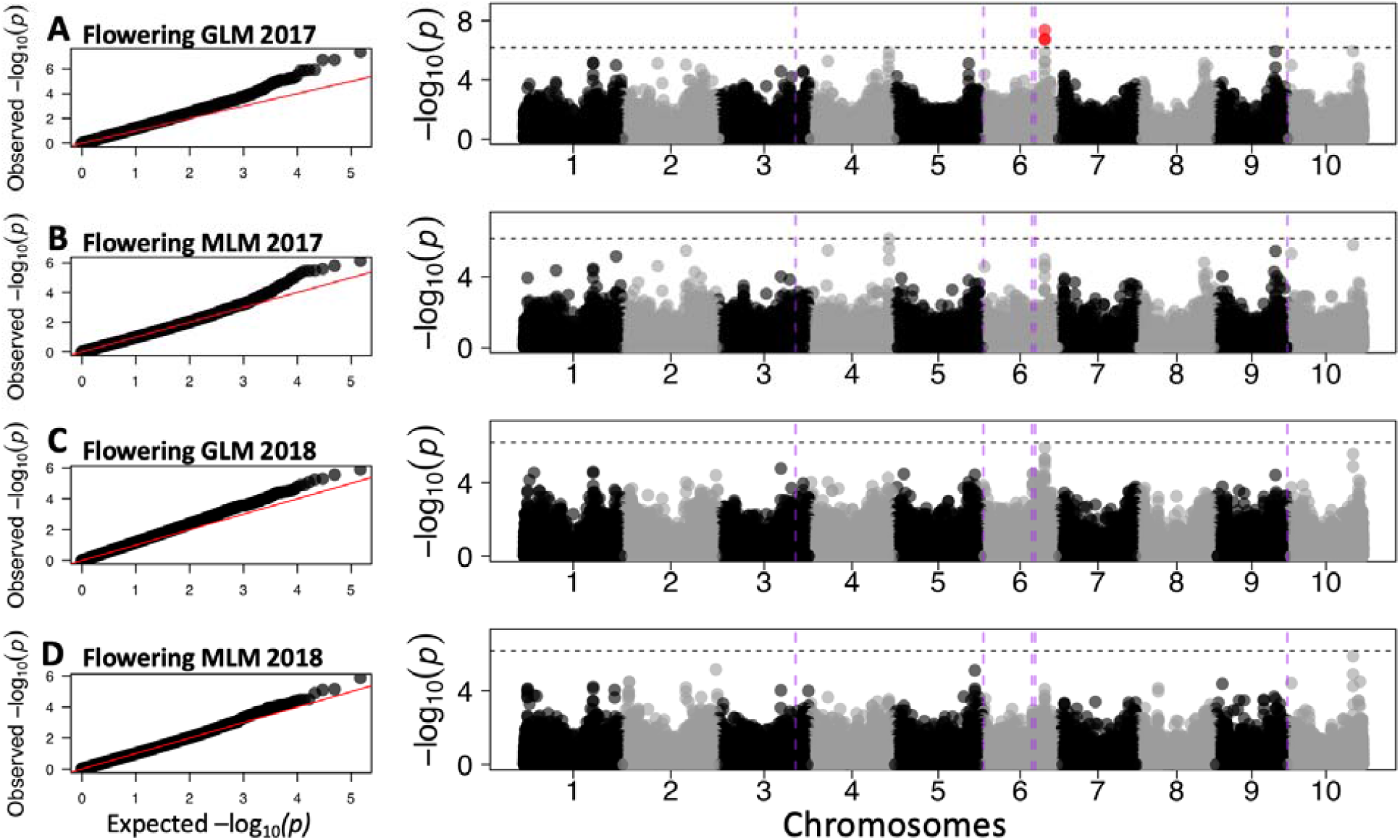
Genome-wide association studies of flowering time in well-watered treatment. (A-B) Quantile-quantile and Manhattan plots in 2017 using the general linear model (GLM) (A) and mixed linear model (MLM) (B). (C-D) Quantile-quantile and Manhattan plots in 2018 using general linear model (GLM) (C) and mixed linear model (MLM) (D). Vertical dashed lines (purple) represent a priori candidate genes for maturity near regions of high association: *SbCN12* (chromosome 3), *Ma6* (chromosome 6), *SbCN4* (chromosome 6), *Ma1* (chromosome 6), and *SbFL9*.*1* (chromosome 9). Horizontal dashed lines represent the threshold at Bonferroni correction (α = 0.05).

**Fig. S7.**
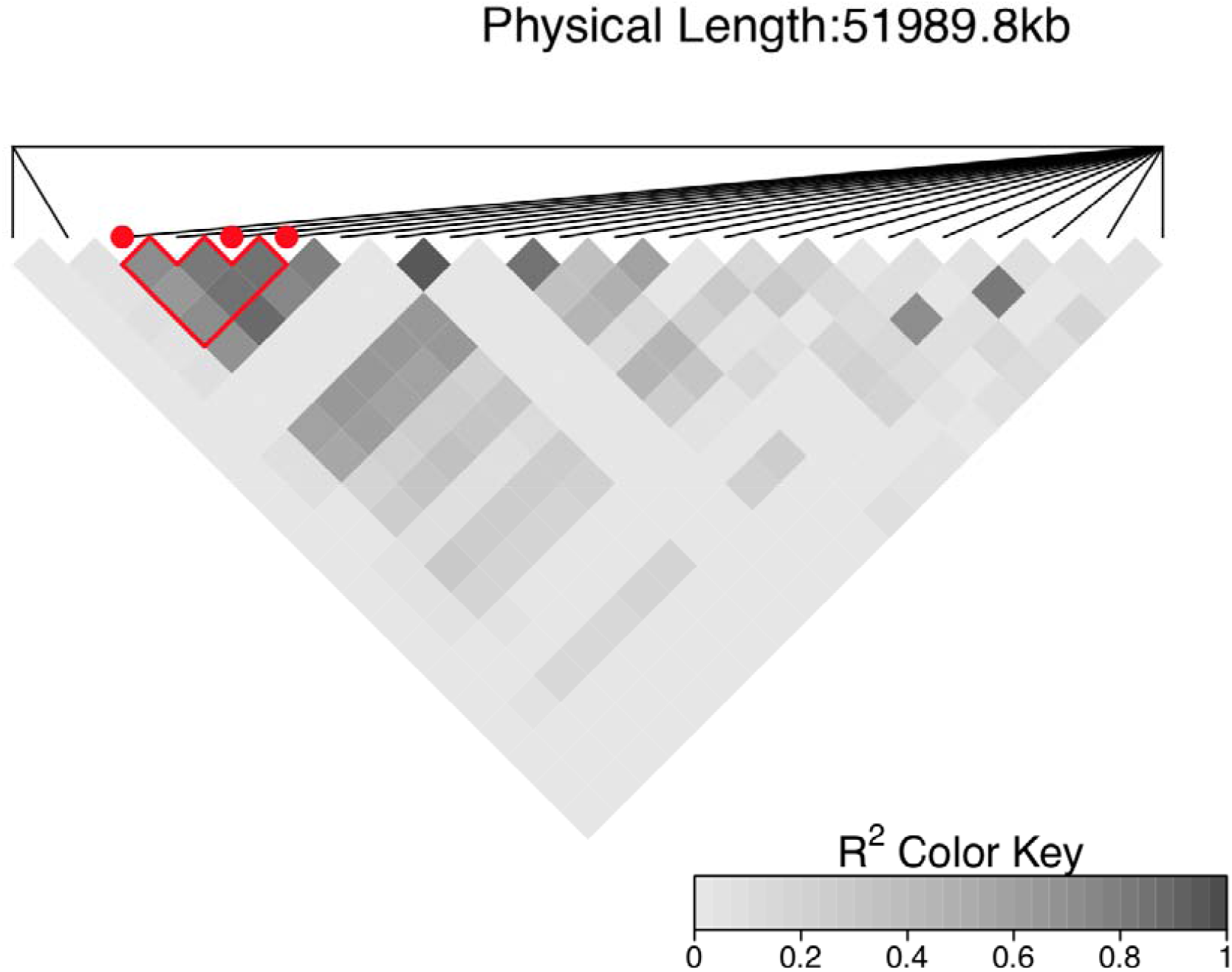
Linkage disequilibrium for around *Stg3a* and trait-associated marker for vegetative biomass under water-stressed treatment. Pairwise R^2^ based heatmap on chromosome 2 for the MTAs S2_57620549, S2_57664163, and S2_57663973 represented in red points. The region selected, within the stay-green gene *Stg3a* (57,146,102 Mb -–61,786,369 Mb), is composed of 22 SNPs including the MTAs for vegetative biomass under water-stressed treatment.

**Table S1.** Country of origin and botanical type of the WASAP_Lysi diversity panel.

| Country | Number | Botanical Type<sup>a</sup> |  |  |  |  |  |  |  |  |
| --- | --- | --- | --- | --- | --- | --- | --- | --- | --- | --- |
|  |  | B | C | D | G | GC | DC | DG | DB | U |
| Mali | 31 | 3 | - | - | 3 | - | - | 1 | 1 | 23 |
| Niger | 116 | 12 | - | 5 | 4 | 21 | 54 | 1 | 3 | 12 |
| Senegal | 37 | - | - | 2 | 29 | 5 | - | 1 | - | - |
| Togo | 28 | - | - | - | - | 3 | - | 1 | - | 24 |
| Other | 7 | - | - | - | - | 4 | - | - | - | 3 |

<sup>a</sup> Botanical type based on morphological classification: B, bicolor; C, caudatum; D, durra; G, guinea; GC, guinea-caudatum; DC, durra-caudatum; DG, durra-guinea; DB, durra-bicolor; U, unknown. "Other" includes international breeding lines.

**Table S2.**
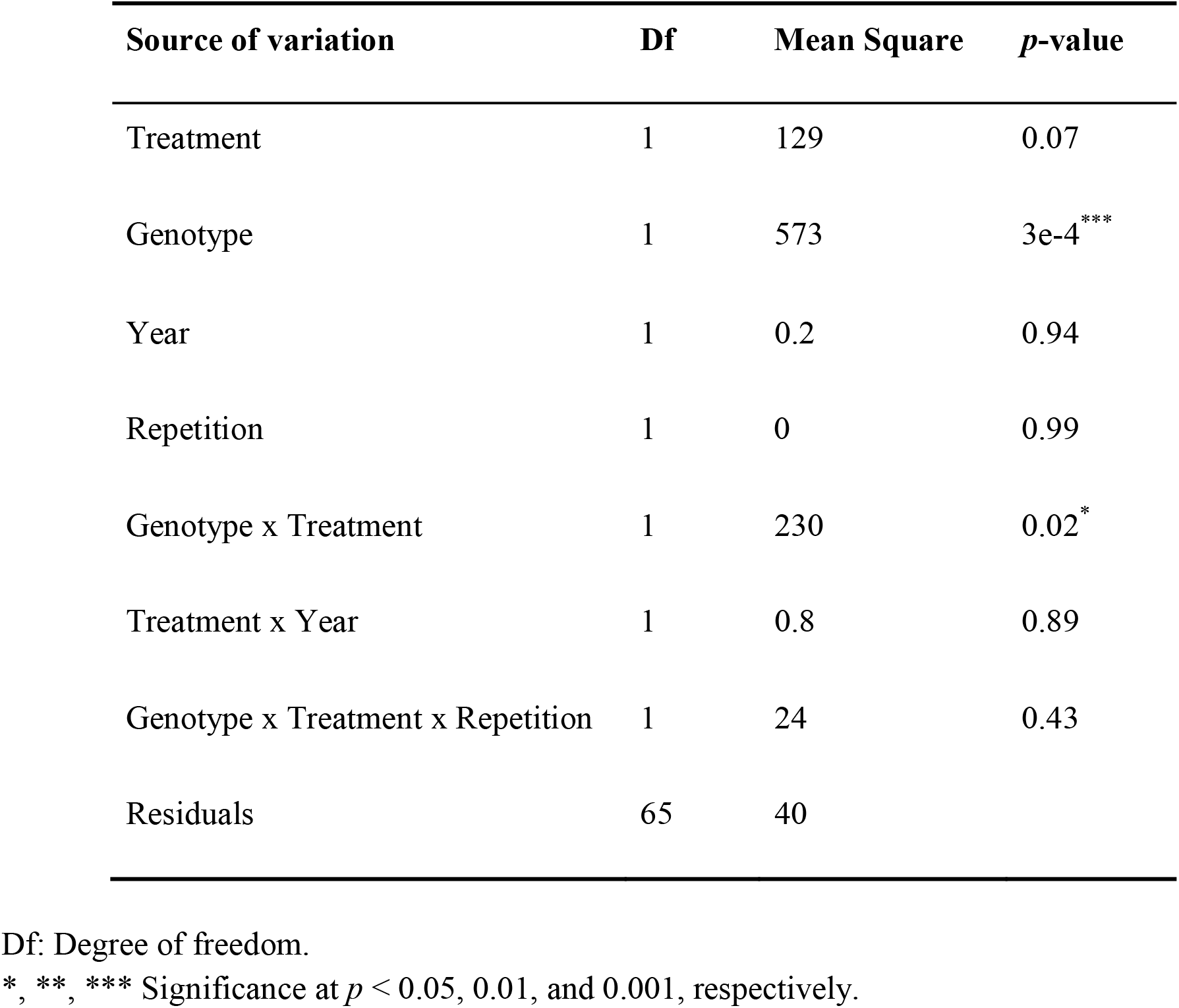
Summary of ANOVA for preflowering drought tolerant genotype (Tx7000) and susceptible genotype (BTx642) for grain weight per panicle for 2017 and 2018.

**Table S3.**
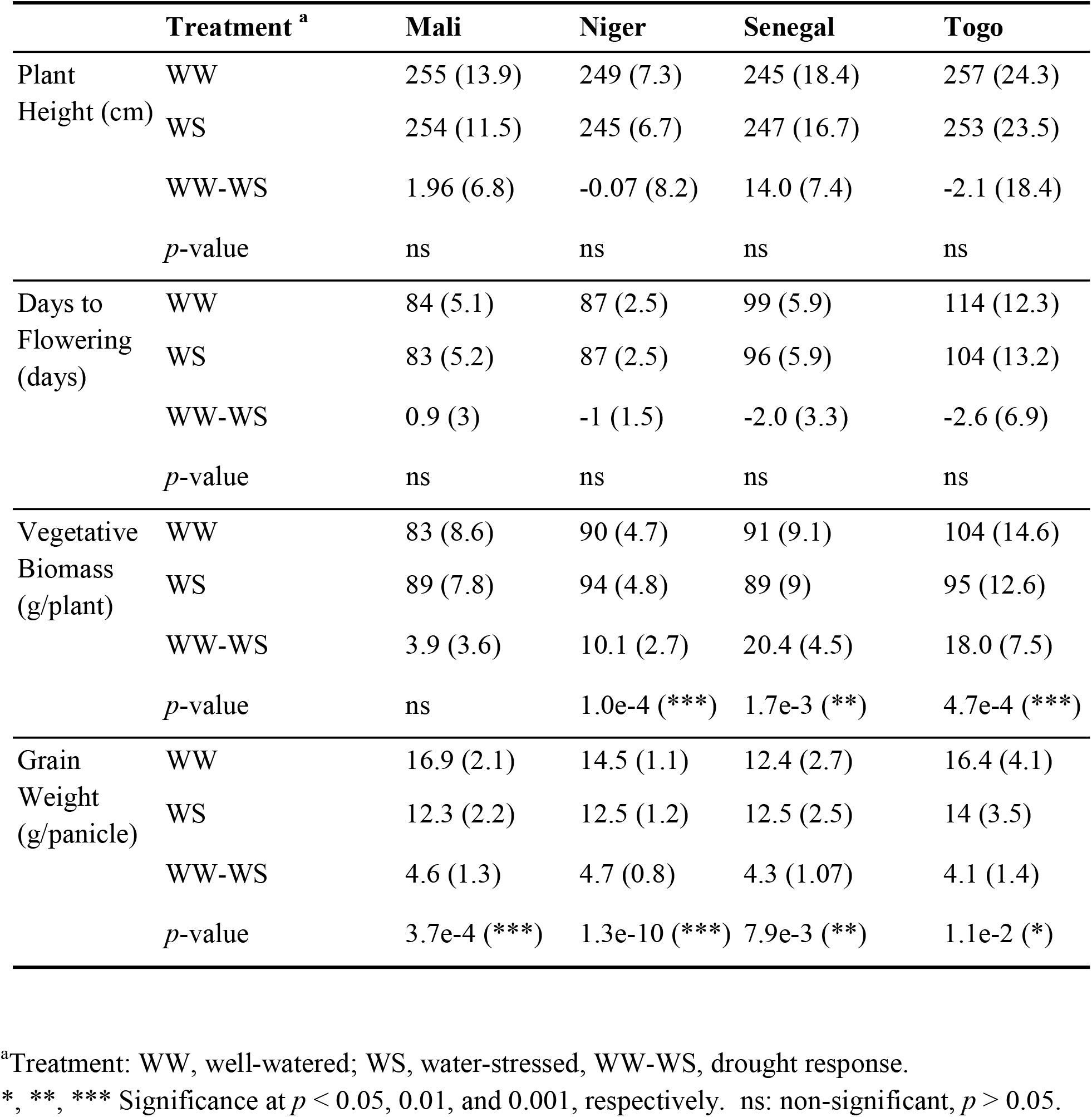
Mean performance (with 95% confidence interval) based on country of origin.

**Table S4.** Mean performance (with 95% confidence interval) based on botanical types.

| **Trait** | **Treatment<sup>a</sup>** | **Caudatum** | **Durra-Caudatum** | **Guinea** |
| --- | --- | --- | --- | --- |
| Plant Height (cm) | WW | 233 (20.5) | 259 (9.1) | 281 (8.4) |
|  | WS | 228 (18.6) | 256 (8.6) | 278 (7.7) |
|  | WW-WS | 5.0 (6) | 0.27 (3.3) | 3.40 (7.78) |
|  | *p*-value | ns | ns | ns |
| Days to Flowering (days) | WW | 89 (5.7) | 81 (6.9) | 105 (8.04) |
|  | WS | 88 (5.5) | 82 (7.1) | 108 (8.6) |
|  | WW-WS | -1.5 (3) | -0.5 (1.4) | -2.5 (2.85) |
|  | *p*-value | ns | ns | ns |
| Vegetative Biomass (g/plant) | WW | 94 (7.8) | 102 (5.6) | 115 (5.4) |
|  | WS | 80 (7.2) | 89 (5.5) | 101 (5) |
|  | WW-WS | 14.7 (3.1) | 9.8 (2.75) | 13.7 (4.1) |
|  | *p*-value | 1e-4 (**) | 1.35e-3 (**) | 3e-4 (***) |
| Grain Weight (g/panicle) | WW | 16.6 (2.2) | 12.5 (1.4) | 9.6 (1.2) |
|  | WS | 10.4 (1.9) | 8.1 (1.2) | 5.7 (1.09) |
|  | WW-WS | 5.6 (1.32) | 4.7 (0.72) | 3.9 (0.67) |
|  | *p*-value | 5.6e-05 (***) | 1.4e-05 (***) | 6.2e-06 (***) |

<sup>a</sup>Treatment: WW, well-watered; WS, water-stressed, WW-WS, drought response.
\*, \*\*, \*\*\* Significance at $p < 0.05$ , 0.01, and 0.001, respectively. ns: non-significant, $p > 0.05$ .

## Supporting Information Files

Data S1: List of genotypes used in the study

Data S2: List of *a priori* candidate genes for maturity and height

Data S3: Marker trait association from the MLMM model

